# Discovery of gene regulatory elements through a new bioinformatics analysis of haploid genetic screens

**DOI:** 10.1101/327775

**Authors:** Bhaven B. Patel, Andres M. Lebensohn, Jan E. Carette, Julia Salzman, Rajat Rohatgi

## Abstract

The systematic identification of regulatory elements that control gene expression remains a challenge. Genetic screens that use untargeted mutagenesis have the potential to identify protein-coding genes, non-coding RNAs and regulatory elements, but their analysis has mainly focused on identifying the former two. To identify regulatory elements, we conducted a new bioinformatics analysis of insertional mutagenesis screens interrogating WNT signaling in haploid human cells. We searched for specific patterns of retroviral gene trap integrations (used as mutagens in haploid screens) in short genomic intervals overlapping with introns and regions upstream of genes. We uncovered atypical patterns of gene trap insertions that were not predicted to disrupt coding sequences, but caused changes in the expression of two key regulators of WNT signaling, suggesting the presence of cis-regulatory elements. Our methodology extends the scope of haploid genetic screens by enabling the identification of regulatory elements that control gene expression.

## Introduction

An outstanding challenge in genomics is the identification of functional regulatory elements that control spatial and temporal expression of protein-coding genes and non-coding RNAs. The Encyclopedia of DNA Elements (ENCODE) project has the ambitious goal of generating a candidate list of all functional elements in the human genome using sequence features, such as conservation, and biochemical features, such as chromatin accessibility and chromatin modifications [1]. Functional approaches to identify regulatory elements have thus far focused on specific regions of the genome and include massively parallel reporter assays or dense clustered regularly-interspaced short palindromic repeats (CRISPR)-mediated mutagenesis of <1 megabase pair segments around a locus of interest (reviewed in [2]).

In work published recently [3], we conducted a comprehensive set of forward genetic screens in haploid human cells to uncover genes required for signaling through the WNT pathway, which plays central roles in development, stem cell function, and cancer. The power of these screens, which used a quantitative transcriptional reporter as the basis for phenotypic selection, was highlighted by the identification of genes encoding both known and novel components that function at most levels of the WNT pathway, from the cell surface to the nucleus. Our previous analysis focused primarily on annotated protein-coding genes and non-coding RNAs. Since the mutant cell libraries used in these screens were generated through untargeted mutagenesis of the genome with a gene trap (GT)-bearing retrovirus, we wondered whether we could use the datasets generated by these screens to uncover gene regulatory mechanisms that modulate the WNT signaling pathway. While retroviral insertions can happen throughout the genome, they are most predominant around transcriptional start sites (TSS), promoters, and enhancers [4]; hence we focused our analysis of retroviral GT insertions on noncoding regions in genes and immediately upstream of them.

Here we present a new bioinformatics pipeline designed to uncover gene regulatory elements and provide evidence for regulatory regions in the first intron of the gene encoding the transcription factor AP4 (TFAP4), a positive regulator of WNT signaling [3], and in the genomic region upstream of the promoter for the gene encoding the WNT co-receptor LRP6.

## Materials and Methods

### Reagents

The reagents used in this manuscript are described in the Materials and methods of [3] and below.

### Antibodies

#### For immunoblotting

Primary antibodies: rabbit anti-AP4 (TFAP4) serum (1:2000, a gift from Takeshi Egawa [5]); rabbit anti-LRP6 (C5C7) (1:500, Cell Signaling Technologies Cat. # 2560); mouse anti-ACTIN (clone C4) (1:500, EMD Millipore Cat. # MAB1501).

Secondary antibodies: peroxidase AffiniPure goat anti-rabbit IgG (H+L) (1:7500, Jackson ImmunoResearch Laboratories Cat. # 111-035-003); IRDye 800CW donkey anti-rabbit IgG (H+L) (1:10,000, Li-Cor Cat. # 925-32213); IRDye 800CW donkey anti-mouse IgG (H+L) (1:10,000, Li-Cor Cat. # 926-32212).

Primary and secondary antibodies used for detection with the Li-Cor Odyssey imaging system were diluted in a 1 to 1 mixture of Odyssey Blocking Buffer (Li-Cor Cat. # 927-40000) and TBST (Tris buffered saline (TBS) + 0.1% Tween-20), and those used for detection by chemiluminescence were diluted in TBST + 5% skim milk. Primary antibody incubations were done overnight at 4°C, and secondary antibody incubations were done for 1 hr at room temperature (RT).

#### For immunostaining

Primary antibodies: mouse anti-LRP6 (clone A59) (5μg/mL, Millipore Cat. # MABS274).

Secondary antibodies: donkey anti-mouse IgG (H+L) Alexa Fluor 647 conjugate (1:200, Thermo 784 Fisher Scientific Cat. # A-31571).

### Cell lines and growth conditions

WT HAP1-7TGP cells and genetically modified clonal derivatives were grown as described in the “Cell lines and growth conditions” section of Materials and methods in [3].

### Bioinformatics analysis

#### Bin-based Analysis of Insertional Mutagenesis Screens (BAIMS)

Genetic screens were conducted as described in the “Reporter-based forward genetic screens” section of Materials and methods in [3], except that GT integrations were mapped as follows. FASTQ files containing 36 bp sequencing reads corresponding to genomic sequences flanking retroviral integration sites in both the sorted and unsorted control cells were obtained for the various genetic screens described (National Center for Biotechnology Information (NCBI) Sequence Read Archive (SRA) Study accession number SRP094861). Reads were aligned to the human genome version “GRCh38” using Bowtie alignment software, version 1.0.1 [6], allowing up to 3 base pair mismatches and only reads that aligned to a single locus of the human genome were considered for downstream analysis. The orientation of the reads relative to the “+” or “−” strand of the chromosome, as defined in human genome version GRCh38, was noted.

Next, the genome was divided into contiguous, non-overlapping intervals of arbitrary length (250–1000 bp as indicated in the Results and figure legends), which are referred to as “bins”, regardless of the location of genes and other genetic elements. Each bin was annotated with any overlapping genes and corresponding features (5’UTR, CDS, intron, and 3’UTR), according to the RefSeq annotations from the University of California, Santa Cruz Table Browser [7] for the GRCh38 assembly of the human genome. An additional genetic feature that we defined as “promoter,” encompassing the 2000 bp directly upstream of the TSS of every gene, was also used to annotate any overlapping bins. The orientation of each genetic feature with respect to the chromosome (whether it resides on the “+” or “−” strand of the chromosome, as specified by the RefSeq annotation) was also noted.

Each GT insertion considered for downstream analysis was mapped to the bin that encompassed its location in the genome. For each bin, we tallied the number of insertions that mapped to the “+” and to the “−” chromosome strand. This enabled us to determine the number of sense and antisense insertions relative to any genetic feature. For example, a GT insertion that aligned to the “+” chromosome strand was considered to be in the sense orientation with respect to a genetic feature that resided on the “+” chromosome strand, whereas an insertion that aligned to the “−” chromosome strand was considered to be in the antisense orientation with respect to the same genetic feature. Histograms depicting the orientation of insertions across genomic regions or genes of interest could then be generated using insertion counts from the bins contained within the region of interest.

#### Gene-based insertion enrichment analysis

To determine which genes were enriched for total GT insertions in the sorted versus the unsorted cells, all insertions in bins annotated with a given gene and its associated promoter as defined above were aggregated separately for the sorted and unsorted cell populations. Thus, the sum of insertions for a specific gene included both sense and antisense insertions that overlapped with the gene’s features, including the promoter. For each gene, a *p*-value for the significance of enrichment was calculated using a one-sided Fisher’s exact test run using the “scipy” package (version 0.7.2) in Python 2.7.5 by comparing the frequency of insertions in the gene in the sorted cells to the frequency of insertions in the gene in the unsorted cells; this *p*-value was then corrected for false-discovery rate. Genes were ranked in ascending order based on FDR-corrected *p*-value.

#### Antisense intronic insertion enrichment analysis

This analysis included bins annotated exclusively as intron and containing at least one GT insertion in the antisense orientation with respect to the gene in the sorted cells. An FDR-corrected *p*-value for the significance of antisense insertion enrichment in each of these bins was determined using a one-sided Fisher’s exact test (from the “scipy” package for Python) comparing the frequency of antisense insertions in the bin for the sorted versus the unsorted cells. Bins were then ranked in ascending order based FDR-corrected *p*-value (Figs 2B and 2C, S1 File).

#### Upstream insertion enrichment analysis

This analysis included bins annotated exclusively as promoter and containing at least one GT insertion regardless of orientation in the sorted cells. An FDR-corrected *p*-value for the significance of insertion enrichment in each of these bins was determined using a one-sided Fisher’s exact test (from “scipy” package for Python) comparing the frequency of insertions in the bin for the sorted versus the unsorted cells. Bins were then ranked in ascending order based on FDR-corrected *p*-value (Figs 2D and 2E, S1 File).

#### Inactivating insertion enrichment analysis

This analysis included bins annotated with any exonic feature (5’UTR, CDS, 3’UTR) and containing at least one GT insertion regardless of orientation in the sorted cells, as well as bins annotated exclusively with intron and containing at least one GT insertion in the sense orientation with respect to the gene in the sorted cells. An FDR-corrected *p*-value for the significance of inactivating insertion (all insertions in bins annotated with 5’UTR, CDS, or 3’UTR and only sense insertions in bins annotated exclusively with intron) enrichment in the bin was determined using a one-sided Fisher’s exact test (from “scipy” package for Python) comparing the frequency of insertions in the bin for the sorted versus the unsorted cells. Bins were ranked in ascending order based on FDR-corrected *p*-value (Figs 2F and 2G, S1 File).

#### BAIMS pipeline code

The BAIMS pipeline code used for the bioinformatics analysis is available through Github (https://github.com/RohatgiLab/BAIMS-Pipeline).

### Isolation of cell lines containing GT insertions

All clonal cell lines containing specified GT insertions were isolated as described in the “Isolation of APC^KO-2^ mutant cell line containing a GT insertion” section of Materials and methods in [3].

The TFAP4^GT^ cell line containing an antisense GT insertion in the first intron of *TFAP4* was isolated from the WNT positive regulator high stringency screen using the reverse primer Wntlow TFAP4 AS II (5’-GCTGCACACGTGTAGACACTC-3’).

LRP6^GT^-1(Up) and LRP6^GT^-2(Up) cell lines, containing antisense GT insertions upstream of the *LRP6* TSS, and the LRP6^GT^-3(Int) cell line, containing a sense GT insertion in the first intron of *LRP6*, were isolated from the WNT positive regulator high stringency screen using the reverse primers LRP6UP-ASGT-Loc-2 (5’-GCAGTGTGTAATATCTCATTCCC-3’), LRP6UP-ASGT-Loc-1 (5’-GGAGACTCCCATTACTCTCTGTT-3’) and Wntlow LRP6 (5’-TGTGGGAAAACTTTGTAATATGC-3’), respectively.

The genomic location of the GT insertion in each isolated cell line is indicated in S5 File.

### Analysis of WNT reporter fluorescence

WNT reporter fluorescence (Figs 3B and 4C) was measured in WT HAP1-7TGP cells or derivatives thereof as described in the “Analysis of WNT reporter fluorescence” section of Materials and methods of [3].

### Quantitative RT-PCR (qPCR) analysis

All mRNA measurements were made as described in the “Quantitative RT-PCR analysis” section of Materials and methods in [3] using the *AXIN2* and *HPRT1* primers described therein (Figs 3C, 3D, 4D, and 4G), the following forward and reverse primers for *TFAP4* (Fig 3D): hTFAP4-RT-PCR-1-FOR (5’-GAGGGCTCTGTAGCCTTGC-3’) and hTFAP4-RT-PCR-1-REV (5’-GAATCCCGCGTTGATGCTCT-3’), and the following forward and reverse primers spanning two pairs of contiguous exons for *LRP6* (Fig 4G): qPCR-LRP6-Exons-1-2-For (5’-GCTTCTGTGTGCTCCTGAG-3’), qPCR-LRP6-Exons-1-2-Rev (5’-TCCAAGCCTCCAACTACAATC-3’), qPCR-LRP6-Exons-7-8-For (5’-GGAGATGCCAAAACAGACAAG-3’), and qPCR-LRP6-Exons-7-8-Rev (5’-CAGTCCAGTAAACATAGTCACCC-3’).

### Immunoblot analysis of WT HAP1-7TGP and mutant cell lines

#### Immunoblot analysis of TFAP4 (Fig 3E)

This analysis was performed as described in the “Immunoblot analysis of HAP1-7TGP and mutant cell lines - Immunoblot analysis of total AXIN1 and AXIN2” section of Materials and methods in [3] with some modifications. Cell pellets harvested from confluent 6 cm dishes were resuspended in 100 μl of ice-cold RIPA lysis buffer (50 mM Tris-HCl pH 8.0, 150 mM NaCl, 2% NP-40, 0.25% deoxycholate, 0.1% SDS, 1X SIGMA*FAST* protease inhibitors, 10% glycerol), sonicated in a Bioruptor 300 (Diagenode) 2 × 30 sec in the high setting, centrifuged 10 min at 20,000 x g, and the supernatant was recovered.

The protein concentration in the supernatant was quantified using the Pierce BCA Protein Assay Kit. Samples were normalized by dilution with RIPA lysis buffer, further diluted with 4X LDS sample buffer supplemented with 50 mM TCEP, heated for 5 min at 95°C, and 40 μg of total protein were electrophoresed alongside Precision Plus Protein All Blue Prestained Protein Standards in NuPAGE 4–12% Bis-Tris gels using 1X NuPAGE MOPS SDS running buffer.

Proteins were transferred to nitrocellulose membranes using 1X NuPAGE transfer buffer + 10% methanol, membranes were stained with Ponceau S to assess loading, washed and blocked with TBST + 5% skim milk. Blots were incubated with rabbit anti-AP4 (TFAP4), washed with TBST, incubated with Peroxidase AffiniPure anti-rabbit secondary antibody, washed with TBST followed by TBS, and developed with SuperSignal™ West Pico Chemiluminescent Substrate and SuperSignal West Femto Maximum Sensitivity Substrate (Thermo Fisher Scientific Cat. # 34080 and 34095).

#### Immunoblot analysis of total LRP6 (Fig 4E and C in S4 Fig)

This analysis was performed as described in the in the previous section. 75 μg of total protein were loaded in duplicate and electrophoresed in a NuPAGE 4–12% Bis-Tris gel. Following the transfer step, the nitrocellulose membrane was cut between the 50 and 75 kDa molecular weight standards and blocked for 1 hour with Odyssey Blocking Buffer. The top membrane was incubated with rabbit anti-LRP6 primary antibody, and the bottom membrane was incubated with mouse anti-ACTIN primary antibody. Membranes were washed with TBST and incubated with IRDye 800CW donkey anti-rabbit IgG and IRDye 800CW donkey anti-mouse IgG secondary antibodies, respectively. Membranes were washed with TBST followed by TBS, and imaged using the Li-Cor Odyssey imaging system. Acquisition parameters in the manufacturer’s Li-Cor Odyssey Image Studio software were set so as to avoid saturated pixels in the bands of interest, and bands were quantified using manual background subtraction. The integrated intensity for LRP6 was normalized to that for ACTIN in the same lane and the average +/− SD from duplicate lanes was used to represent the data in Fig 4E.

### LRP6 cell-surface staining of WT HAP1-7TGP and mutant cell lines (Fig 4F)

Approximately 24 hr before staining, cells were seeded in a 6-well plate at a density of 2×10^5^ per well and grown in 2.5 ml of CGM 2 (defined in [3]). On the following day the cells were washed once in 3 ml of phosphate buffered saline (PBS), harvested in 0.5 ml of Accutase Cell Detachment Reagent (Thermo Fisher Scientific Cat. # NC9839010), resuspended in 1.5 ml of ice-cold CGM 2 and centrifuged 4 min at 400 × g at 4°C (all subsequent centrifugation steps were done in the same way). Cells were washed with 2 ml of ice-cold Iscove’s Modified Dulbecco’s Medium (IMDM) with L-glutamine, with HEPES, without Alpha-Thioglycerol (GE Healthcare Life Sciences Cat. # SH30228.01) + 1% Fetal Bovine Serum (FBS) (Atlanta Biologicals Cat. # S11150), centrifuged and resuspended in 150 μl of mouse anti-LRP6 primary antibody in IMDM + 1% FBS. Following a 1 hr incubation on ice, cells were washed with 1.8 ml of ice-cold IMDM + 1% FBS, centrifuged, washed with 2 ml of ice-cold IMDM +1% FBS and centrifuged again. Cells were resuspended in 150 μl of secondary antibody in ice-cold IMDM + 1% FBS and incubated on ice for 30 minutes. Cells were washed with 1.8 ml of ice-cold IMDM + 1% FBS, centrifuged, washed with 2 ml of ice-cold IMDM + 1% FBS and centrifuged again. Cells were resuspended in 200 μl of PBS + 2% FBS and LRP6 cell-surface fluorescence was analyzed on a BD Accuri RUO Special Order System (BD Biosciences).

## Results

### Bin-based Analysis of Insertional Mutagenesis Screens (BAIMS)

Haploid genetic screens rely on the phenotypic selection of a population of cells mutagenized by integration of a GT-bearing retrovirus. GTs, which contain a splice acceptor (SA) and a transcriptional termination polyadenylation signal (pA), can disrupt protein-coding genes in two ways: (1) by inserting into an exon in either the sense or antisense orientation relative to the coding sequence of the gene or (2) by inserting into an intron in the sense orientation, such that the directional SA causes the GT to be spliced to the 3’-end of the preceding exon, resulting in a transcript that undergoes premature termination (A-D in S2 Fig). Indeed, most hits in haploid genetic screens exhibit a bias towards such inactivating sense insertions in introns [8]. GT insertions can theoretically also perturb gene regulation by directly interrupting a regulatory protein-binding site on DNA, by terminating a regulatory transcript, or by altering chromatin structure. Therefore, in principle it should be possible to find GT insertion patterns indicative of such regulatory mechanisms.

In order to map GT insertions in a way that would enable us to identify regulatory elements, we devised a bioinformatics pipeline that was completely agnostic to the boundaries of annotated genes and simply tracked the number and orientation of GT insertions in short genomic intervals of arbitrary size, defined as “bins” (Fig 1A). We refer to this approach as “<u>B</u>in-based <u>A</u>nalysis of <u>I</u>nsertional <u>M</u>utagenesis <u>S</u>creens”, or BAIMS. Sequencing reads adjacent to the location of GT insertions found in sorted (phenotypically selected) and unsorted (control) cells from haploid genetic screens were aligned to the human genome and assigned to the bin that encompassed the insertion (Fig 1B). The orientation of each insertion on the chromosome was defined according to the GRCh38 assembly of the human genome. Each bin was also annotated with any relevant genetic features it overlapped with (5’ untranslated region (5’UTR), coding domain sequence (CDS), intron and 3’ untranslated region (3’UTR)), using the RefSeq annotations from the University of California, Santa Cruz Table Browser [7] for the GRCh38 assembly of the human genome. We also defined an additional genetic feature, designated “promoter”, as the 2000 base pairs (bp) upstream of the TSS of each gene. This region typically includes the minimal promoter but may also contain other cis-regulatory elements. We annotated bins overlapping with this feature accordingly. The relative orientation of any insertion with respect to any feature can therefore be determined, allowing us to observe patterns of sense and antisense GT insertions across features of interest (Fig 1C). This information can be displayed in a histogram depicting insertions over any genomic region of interest (Fig 1C), providing a high-resolution picture of the insertional landscape. Thus, BAIMS enables us to identify atypical patterns of GT insertions in specific genetic features that could be indicative of regulatory elements.

**Fig 1.**
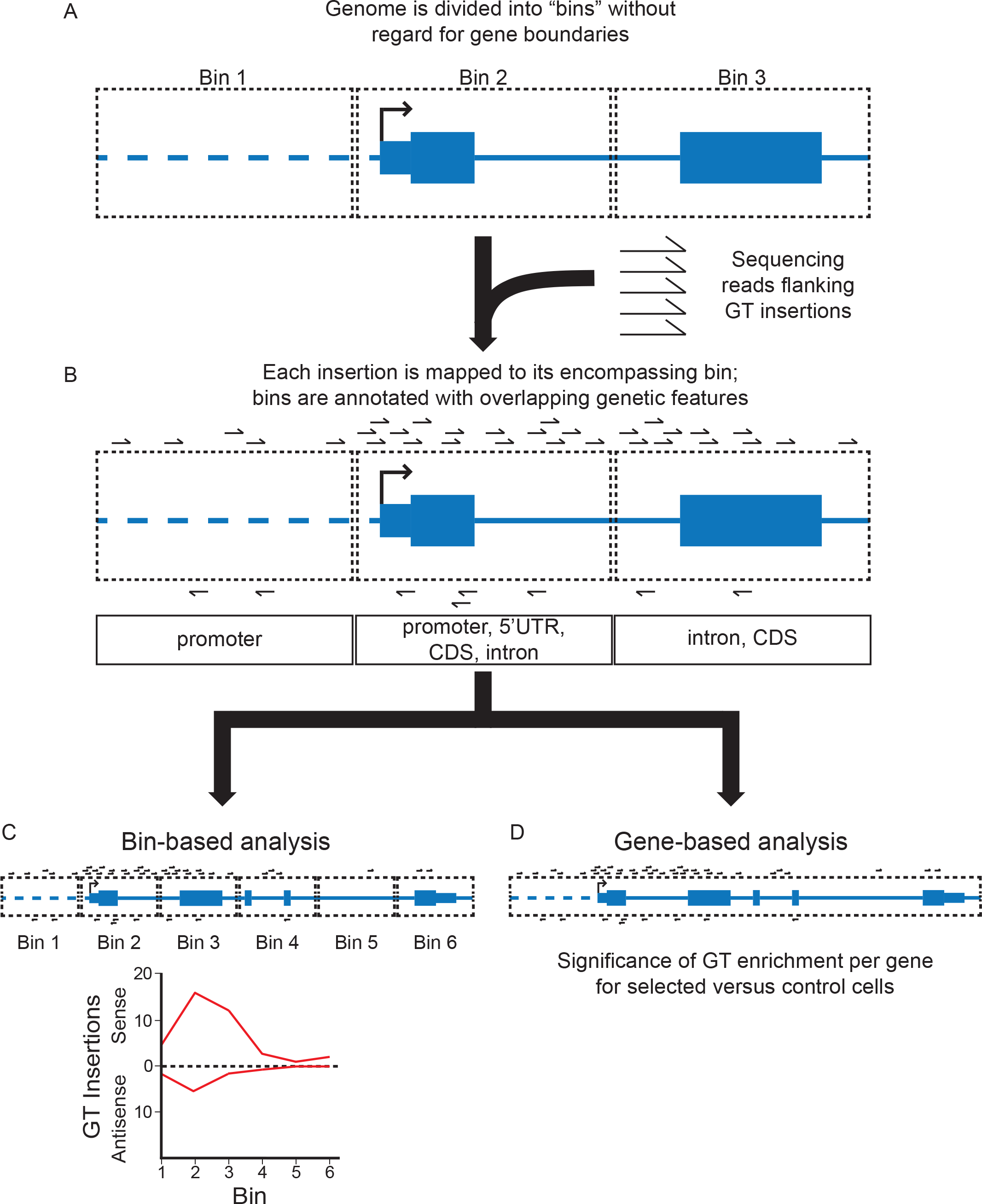
Schematic of Bin-based Analysis of Insertional Mutagenesis Screens (BAIMS). (A) The human genome is computationally divided into “bins” (pictured as rectangles with black dotted lines), which encompass contiguous segments of DNA of an equal arbitrary length. Throughout this study, we used bins of 250 bp or 1000 bp in length, depending on the resolution required for the analysis. The boundaries of annotated genetic features, including genes and regulatory elements, are ignored. The depicted fictitious gene is modeled after a RefSeq gene track following the University of California, Santa Cruz (UCSC) genome browser display conventions: coding exons are represented by tall blocks, UTRs by shorter blocks, and introns by horizontal lines connecting the blocks. The arrow indicates the gene’s TSS. (B) Sequencing reads flanking the location of individual GT insertions in the control and selected cell populations from a haploid genetic screen are mapped to the human genome and assigned to the bin that encompasses the location of the insertion. The orientation of each insertion relative to the chromosome is noted. Bins are also annotated with any overlapping genetic features. These include promoter (defined as the 2000 bp upstream of the TSS, indicated by a horizontal dotted line), 5’UTR, CDS, intron, and 3’UTR. The orientation of the feature relative to the chromosome is also noted. (C) For the bin-based analysis, the number and orientation of GT insertions in consecutive bins along any defined portion of the genome (including but not limited to genes) is determined and can be depicted in a histogram (the number of sense GT insertions per bin is arbitrarily shown above the horizontal line labeled “0”, and the number of antisense insertions below), enabling the visualization of insertion patterns at sub-gene resolution. (D) For the gene-based analysis, GT insertions in bins that overlap with genes can be summed to obtain a total insertion count for each gene. The significance of GT enrichment for every gene is calculated by comparing the total number of insertions per gene found in the selected versus the control cell populations (see Materials and Methods for details).

The overall enrichment of GT insertions for any given gene in the selected versus the control cells from a haploid genetic screen can also be assessed by aggregating the insertions found in all bins that overlap with the gene (Fig 1D; see Materials and Methods). We refer to this analysis, which produces a significance score for GT enrichment comparable to that of previous analyses [3], as “gene-based insertion enrichment analysis”.

### Identification of regulatory elements through the analysis of bins with atypical GT insertion patterns

Our previous analysis [3] focused on GT insertions predicted to inactivate protein-coding genes and non-coding RNAs as outlined above: sense and antisense insertions in exons, and sense insertions in introns (B-D in S2 Fig). To identify regulatory elements, we searched for GT insertion patterns distinct from these. Because the GT retrovirus has a strong propensity to integrate near TSSs, promoters and enhancers, we limited our analysis to non-coding regions within and adjacent to genes. We used BAIMS to look for enrichment of antisense insertions in introns, which would not be predicted to interrupt protein-coding transcripts (E in S2 Fig), and for enrichment of insertions in either orientation in the regions upstream of the TSS of genes (F and G in S2 Fig). Since each bin is annotated with the genetic features it overlaps with (Fig 2A), we could readily identify these distinct patterns of GT insertions.

**Fig 2.**
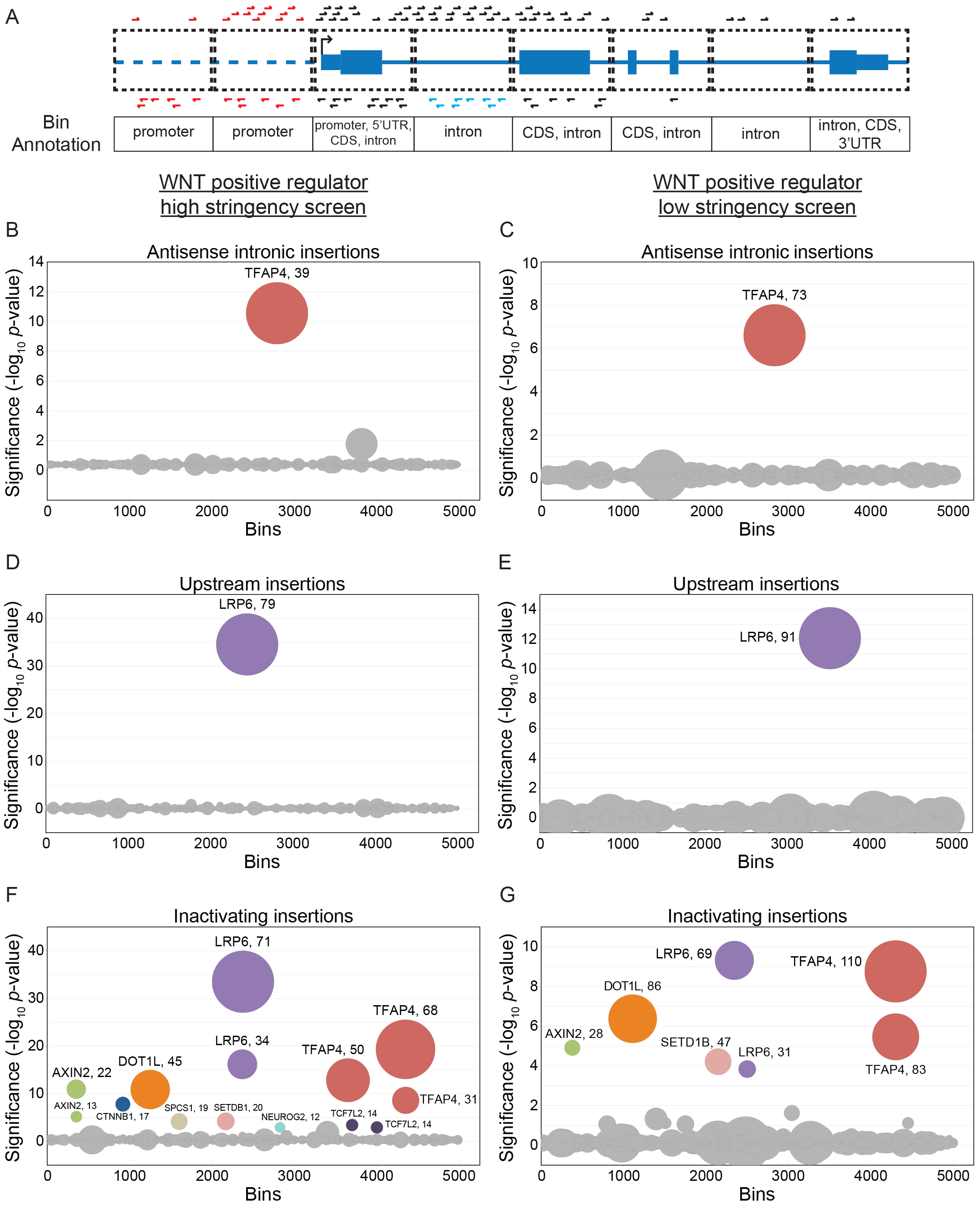
BAIMS identifies atypical GT insertion patterns in screens for regulators of WNT signaling. (A) Schematic depicting various patterns of GT insertions relative to genetic features in the containing bins, used for the antisense intronic, upstream and inactivating insertion enrichment analyses (see text for details). A fictitious gene modeled after a RefSeq gene track, with GT insertions in the sense orientation relative to the gene depicted above the track and in the antisense orientation depicted below it. The antisense intronic insertion enrichment analysis accounts for antisense GT insertions in bins annotated exclusively as intron (depicted in blue) and the upstream insertion enrichment analysis accounts for both sense and antisense insertions in bins annotated exclusively as promoter (depicted in red). These two classes of insertions had been ignored in previous gene-based analyses of haploid genetic screens [3]. The inactivating insertion enrichment analysis accounts for both sense and antisense insertions in bins annotated as 5’UTR, CDS, or 3’UTR, as well as sense insertions in bins annotated exclusively as intron; these insertions (depicted in black) include all the gene-inactivating insertions used in previous analyses. (B-G) Circle plots depicting the results of antisense intronic (B, C), upstream (D, E), and inactivating (F, G) insertion enrichment analyses for the WNT positive regulator high stringency (B, D, and F) and low stringency (C, E, and G) screens. Circles represent individual 1000 bp bins. The y-axis indicates the significance of GT insertion enrichment in the sorted versus the unsorted, control cells, expressed in units of −log_10_(FDR-corrected *p*-value), and the x-axis indicates the 5000 bins with the smallest FDR-corrected *p*-values, arranged in random order. Circles representing bins with an FDR-corrected *p*-value < 0.01 are colored and labeled with the name of the gene with which the bin overlaps. Circles representing bins corresponding to the same gene are depicted in the same color. The diameter of each circle is proportional to the number of independent GT insertions mapped to the corresponding bin in the sorted cells, which is also indicated next to the gene name for enriched bins.

To identify regulatory elements in introns, we looked for enrichment of antisense insertions in bins exclusively annotated as intron (Fig 2A); we refer to this analysis as “antisense intronic insertion enrichment analysis.” To identify regulatory elements in regions immediately upstream of genes, we looked for enrichment of both sense and antisense GT insertions in bins exclusively annotated as promoter (Fig 2A); we refer to this analysis as “upstream insertion enrichment analysis.” To distinguish features identified in the previous two analyses from the more typical disruption of protein-coding genes or non-coding RNAs by GT insertions, we looked for enrichment of gene-inactivating insertions, as defined above (sense and antisense insertions in bins annotated with 5’UTR, CDS or 3’UTR, and sense insertions in bins annotated exclusively as intron; see Fig 2A); we refer to this analysis as “inactivating insertion enrichment analysis.”

These three analyses were applied to the data from two screens for positive regulators of signaling following stimulation with WNT3A, henceforth referred to as the WNT positive regulator high and low stringency screens, which differed only in the stringency of selection [3]. In these screens, HAP1 cells harboring a WNT-responsive GFP reporter (hereafter referred to as “WT HAP1-7TGP”) were mutagenized with GT retrovirus, treated with WNT3A and sorted by fluorescence activated cell sorting (FACS) for cells that exhibited the lowest 2% (high stringency) or the lowest 10% (low stringency) signaling activity. These screens enabled us to identify known and new regulators in the WNT pathway [3].

Antisense intronic insertion enrichment analysis of the WNT positive regulator high and low stringency screens produced only one significant (FDR-corrected *p*-value < 0.01) bin (Figs 2B and 2C, S1 File), which mapped to the gene *TFAP4*, one of the top hits from these screens [3]. Upstream insertion enrichment analysis of the same screens produced only one significant bin upstream of *LRP6* (Figs 2D and 2E, S1 File), which was the top hit of both of these screens [3]. These results are markedly different from those of the inactivating insertion enrichment analysis of the same screens (Figs 2F and 2G, S1 File), which revealed bins in many of the same genes identified as significant hits in these screens [3].

In the sections that follow, we tested if the GT insertion patterns identified in *TFAP4* and *LRP6* by the antisense intronic insertion and upstream insertion enrichment analyses, respectively, reflected regulatory effects on gene expression.

### Antisense GT insertions in the first intron of *TFAP4* disrupt gene expression

The second top hit in the WNT positive regulator high and low stringency screens was *TFAP4*, encoding the transcription factor TFAP4, which we have shown to be a positive regulator of the WNT pathway acting downstream of the key transcriptional co-activator β-catenin (CTNNB1) [3]. As is common for top hits of haploid genetic screens, the 5’ end of *TFAP4* was significantly enriched for inactivating GT insertions, including many sense and antisense insertions in the first exon as well as sense insertions in the first intron, which are all expected to disrupt the *TFAP4* coding sequence (Fig 3A and A in S3 Fig). However, the single bin identified in the antisense intronic insertion enrichment analysis (Figs 2B and 2C, S1 File) was also located in the first intron and it contained a comparable number of antisense GT insertions (Fig 3A and A in S3 Fig), which would not be predicted to disrupt the *TFAP4* coding sequence. This pattern of GT insertion enrichment was not seen for *TFAP4* in the mutagenized but unsorted cells used as a control for the WNT positive regulator screens (Fig 3A) or for other top hits, such as *DOT1L*, in the sorted cells from these same screens (B in S3 Fig). These results suggested that antisense GT insertions in the first intron of *TFAP4* (which would not be predicted to terminate the *TFAP4* transcript) reduced WNT signaling.

**Fig 3.**
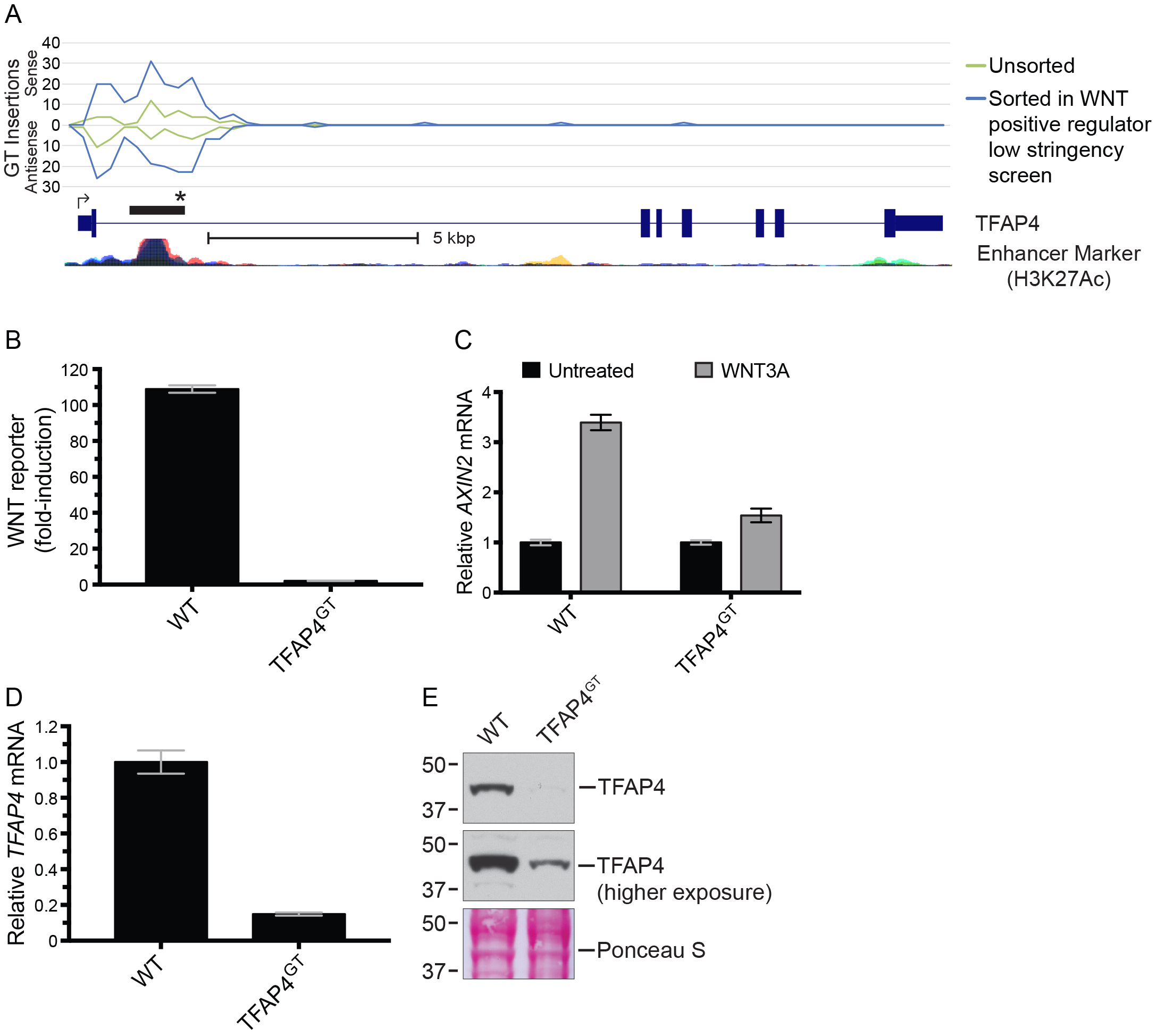
Antisense GT insertions in the first intron of *TFAP4* disrupt gene expression and impair WNT signaling. (A) The histogram indicates the number and orientation of GT insertions mapped to *TFAP4* in unsorted cells and in the sorted cells from the WNT positive regulator low stringency screen. Values above the horizontal line labeled “0” indicate sense insertions relative to the coding sequence of the gene, and values below it indicate antisense insertions. The x-axis represents contiguous 250 bp bins to which insertions were mapped (Chromosome 16, 4257249-4273000 bp). Insertions mapped for the different cell populations indicated in the legend are depicted by traces of different colors. A RefSeq gene track for *TFAP4* (following UCSC genome browser display conventions, described in the legend of Fig 1A) and an ENCODE track for histone3-lysine27-acetylation, a marker for enhancer activity (taken from the UCSC genome browser), are shown underneath the graph. The black rectangle above the gene track indicates the location of the bin identified in the antisense intronic insertion enrichment analyses of both the WNT positive regulator low stringency and high stringency screens. The black star denotes the position of the antisense GT insertion in the TFAP4^GT^ clonal cell line used for further characterization. A scale bar is provided beneath the gene track for reference. (B) Fold-induction in WNT reporter (median +/− standard error of the median (SEM) EGFP fluorescence from 10,000 cells) following treatment with 50% WNT3A conditioned media (CM). (C) *AXIN2* mRNA (average +/− standard deviation (SD) of *AXIN2* mRNA normalized to *HPRT1* mRNA, each measured in triplicate qPCR reactions) relative to untreated cells. Where indicated, cells were treated with 50% WNT3A CM. (D) *TFAP4* mRNA (average +/− SD of *TFAP4* mRNA normalized to *HPRT1* mRNA, each measured in triplicate qPCR reactions) relative to WT HAP1-7TGP cells. (E) Immunoblot of TFAP4. The middle panel shows a higher exposure of the same blot shown in the top panel, and the bottom panel displays Ponceau S staining of the same blot as a loading control. Molecular weight standards in kiloDaltons (kDa) are indicated to the left of each blot.

To confirm this prediction, we isolated a clonal cell line harboring an antisense GT insertion in the first intron of *TFAP4* (we designate this cell line TFAP4^GT^; see Fig 3A and Materials and Methods). WNT3A-induced reporter activation was nearly eliminated in TFAP4^GT^ cells when compared to WT HAP1-7TGP cells (Fig 3B). Expression of *AXIN2* mRNA, a universal target gene of the pathway, following treatment with WNT3A was also reduced substantially in TFAP4^GT^ cells (Fig 3C). Given its location within the boundaries of the *TFAP4* gene, we tested whether the antisense GT insertion affected expression of *TFAP4* itself. Both *TFAP4* mRNA and protein levels were severely reduced in TFAP4^GT^ cells, explaining the observed defect in pathway activity (Figs 3D and 3E). A higher exposure of the TFAP4 immunoblot from TFAP4^GT^ cells revealed a faint band corresponding to TFAP4 (Fig 3E), indicating that the antisense GT insertion in the first intron of *TFAP4* reduced expression of a full-length transcript and protein as opposed to disrupting the coding sequence.

### Antisense GT insertions upstream of *LRP6* reduce LRP6 protein abundance independently of mRNA levels

*LRP6* encodes a required co-receptor for WNT ligands and was the top hit of the WNT positive regulator high and low stringency screens [3]. As expected, most GT insertions in the *LRP6* gene proper (downstream of the TSS) were in the sense orientation with respect to the coding sequence (Figs 4A and 4B, and A and B in S4 Fig). However, our upstream insertion enrichment analysis also revealed a bin enriched for GT insertions located upstream of the TSS (Figs 2D and 2E, S1 File). A closer inspection of the region surrounding this bin revealed a pronounced enrichment of antisense insertions extending from about 1 to 3.5 kilobase pairs (kbp) upstream of the TSS (Fig 4B and B in S4 Fig). Importantly, this region was located upstream of the annotated *LRP6* promoter in *Ensembl* (Fig 4B). These GT insertion patterns were not observed in the mutagenized but unsorted cells used as a control for the WNT positive regulator screens (Figs 4A and 4B). These results suggested that antisense insertions upstream of *LRP6* impaired WNT signaling.

**Fig 4.**
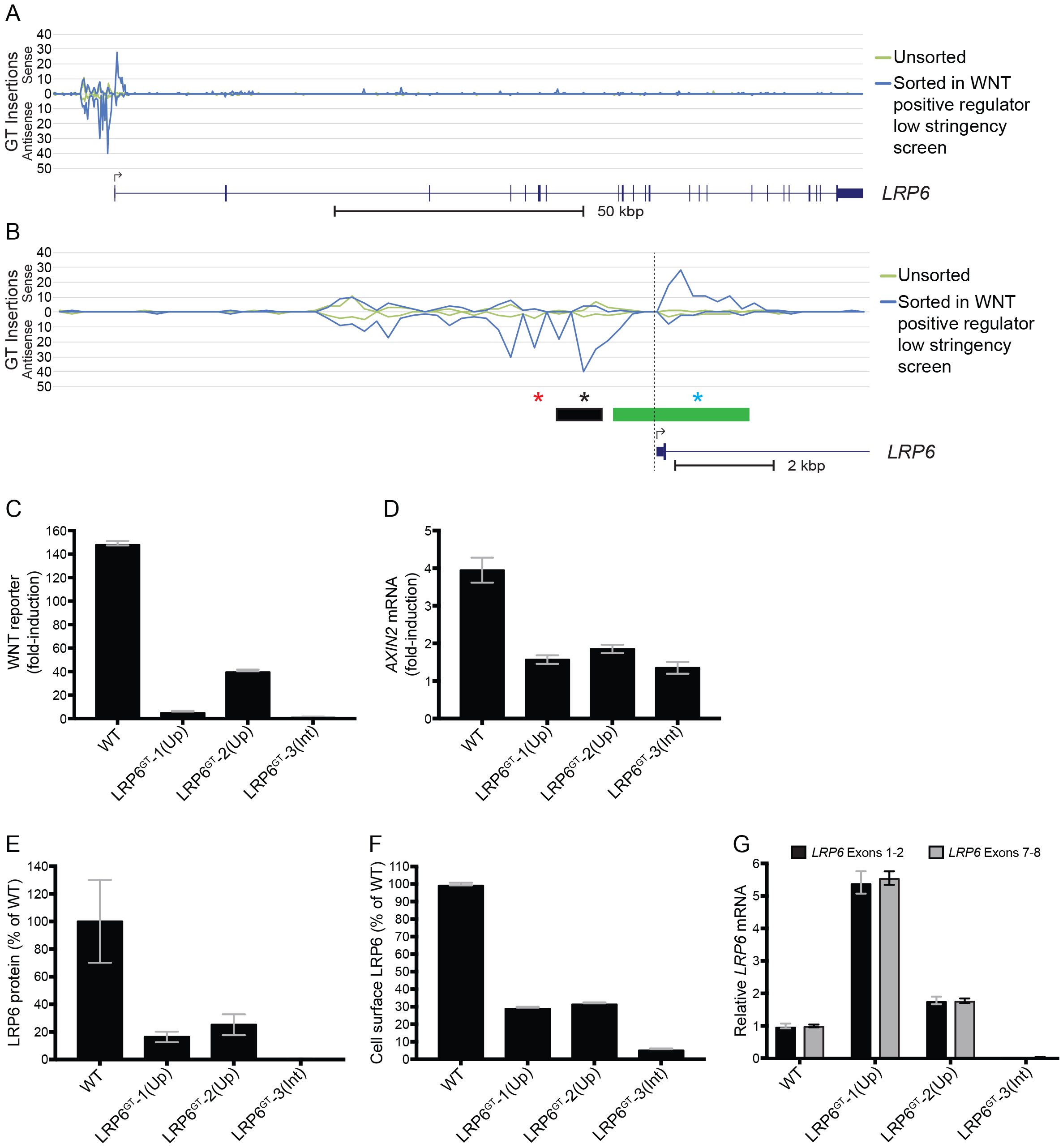
Antisense GT insertions upstream of *LRP6* reduce LRP6 protein expression and impair WNT signaling. (A) The histogram indicates the number and orientation of GT insertions mapped to *LRP6* and to the region ~12.5 kbp upstream of the TSS in unsorted cells and in the sorted cells from the WNT positive regulator low stringency screen. See legend to Fig 3A for details. The x-axis represents contiguous 250 bp bins to which insertions were mapped (Chromosome 12, 12116000-12279249 bp). (B) The histogram shows an expanded view of the 5’ end of *LRP6* and the region ~12.5 kbp upstream of the TSS (left of the vertical dotted line), with traces for GT insertions mapped in unsorted cells and in the sorted cells from the WNT positive regulator low stringency screen. The x-axis represents contiguous 250 bp bins to which insertions were mapped (Chromosome 12, 12262500-12279249 bp). The green rectangle above the gene track indicates the location of the *LRP6* promoter according to *Ensembl* and the black rectangle indicates the location of the bin identified in the upstream insertion enrichment analyses of the WNT positive regulator low stringency and high stringency screens. The black and red stars denote the position of the antisense GT insertions in the LRP6^GT^-1(Up) and LRP6^GT^-2(Up) clonal cell lines, respectively, and the blue star denotes the position of the sense GT insertion in the LRP6^GT^-3(Int) cell line. (C) Fold-induction in WNT reporter (median +/− SEM EGFP fluorescence from 20,000 cells) following treatment with 50% WNT3A CM. (D) Fold-induction in *AXIN2* mRNA (average +/− SD of *AXIN2* mRNA normalized to *HPRT1* mRNA, each measured in triplicate qPCR reactions) following treatment with 50% WNT3A CM. (E) Quantification of immunoblot analysis of total LRP6 protein (average +/− SD LRP6 intensity normalized to ACTIN intensity from samples run in duplicate) shown as percentage of WT HAP1-7TGP. The blot used for quantification is shown in C in S4 Fig. (F) Cell surface LRP6 protein (median +/− SEM cell surface LRP6 immunofluorescence from 20,000 cells) shown as percentage of WT HAP1-7TGP. (G) *LRP6* mRNA (average +/− SD of *LRP6* mRNA, measured using two different primer pairs, normalized to *HPRT1* mRNA, each measured in triplicate qPCR reactions) shown relative to WT HAP1-7TGP cells.

To test this possibility, we isolated two clonal cell lines containing antisense GT insertions in the region upstream of *LRP6* (we designate these cell lines LRP6^GT^-1(Up) and LRP6^GT^-2(Up); see Fig 4B and Materials and Methods) and as a control we isolated a clonal cell line with a sense GT insertion in the first intron of *LRP6* that is predicted to disrupt the *LRP6* coding sequence (we designate this cell line LRP6^GT^-3(Int); see Fig 4B and Materials and Methods). Both LRP6^GT^-1(Up) and LRP6^GT^-2(Up) cells demonstrated significantly reduced WNT reporter activation and *AXIN2* mRNA accumulation following treatment with WNT3A when compared to WT HAP1-7TGP cells (Figs 4C and 4D). The most plausible explanation for how the GT insertions reduced WNT signaling would be down-regulation of LRP6, which is indeed what we observed when we measured total and cell-surface levels of LRP6 protein. LRP6^GT^-1(Up) and LRP6^GT^-2(Up) cells exhibited a 75–84% reduction in total LRP6 protein and a 68–71% reduction in cell-surface LRP6 compared to WT cells (Figs 4E and 4F). LRP6^GT^-3(Int) cells exhibited greater, >99% and 94% reductions in total and cell-surface LRP6, respectively, compared to WT cells, as would be expected from the disruption of the *LRP6* coding sequence caused by the sense GT insertion in the first intron (Figs 4E and 4F).

Unexpectedly, despite the reduction in LRP6 protein observed in LRP6^GT^-1(Up) and LRP6^GT^-2(Up) cells harboring antisense GT insertions upstream of the *LRP6* promoter, we did not observe a corresponding decrease in *LRP6* mRNA (Fig 4G). In an important control, *LRP6* mRNA levels were indeed markedly reduced in LRP6^GT^-3(Int) cells carrying a sense intronic GT insertion that disrupts the coding sequence (Fig 4G). These results suggest that antisense GT insertions upstream of *LRP6* diminished signaling by an enigmatic mechanism that reduced LRP6 protein levels without altering mRNA levels, rather than by simply disrupting the *LRP6* promoter. Interestingly, sequence elements with similar properties have been described upstream of promoter elements for heat shock target genes in yeast [9].

## Discussion

We developed a new bioinformatics tool to analyze haploid genetic screens with the explicit goal of uncovering regulatory elements. We analyzed screen data in a way that discerned GT insertion patterns distinct from those predicted to disrupt the coding sequence of genes, and found that atypical insertions in introns and regions upstream of the TSS can cause down-regulation of genes, leading to the phenotype selected for during the screen. When we applied this new analysis to haploid genetic screens interrogating the WNT pathway, we found that antisense GT insertions in the first intron of *TFAP4* and upstream of the *LRP6* promoter resulted in marked changes in the expression of these genes. These types of insertions had not been accounted for in previous analyses of haploid genetic screens.

The identified GT insertions could disrupt regulatory elements such as promoters, enhancers, antisense transcripts or splicing sequences. In the case of *TFAP4*, most of the insertions were located in the first intron and overlapped with a strong enhancer signal (Fig 3A), suggesting they may disrupt an enhancer. Previous studies have shown that *TFAP4* is directly regulated by c-MYC and that the first intron of *TFAP4* in fact contains four c-MYC binding sites [10, 11], three of which are encompassed by the bin identified in our antisense intronic insertion enrichment analysis (Figs 2B and 2C). In future studies, it will be important to test whether the antisense insertions found in the first intron of *TFAP4* down-regulate *TFAP4* mRNA (Fig 3D) by disrupting c-MYC binding or through an alternative mechanism.

Similarly, LRP6 protein was down-regulated in the LRP6^GT^-1(Up) and LRP6^GT^-2(Up) cell lines containing antisense GT insertions upstream of the *LRP6* promoter (Figs 4E and 4F). Surprisingly, *LRP6* mRNA levels were not reduced in these same cell lines, suggesting a mechanism regulating LRP6 protein levels. In yeast, genomic sequences upstream of genes that have no effect on mRNA levels can instead regulate protein levels [9]. The selective enrichment of antisense versus sense GT insertions in the region upstream of the *LRP6* promoter in cells sorted for low WNT reporter fluorescence (Figs 4A and 4B) suggests that such insertions are not merely disrupting an enhancer or promoter. Instead, we speculate that these GT insertions may disrupt an antisense transcript or another directional element residing on the antisense strand that positively regulates *LRP6* expression. Since no such elements have been described, it will be important to elucidate the nature of this regulatory mechanism in future studies.

Unlike other more focused efforts to identify regulatory regions associated with a given gene or set of genes [12–18], our untargeted approach enables the genome-wide identification of cis-regulatory elements involved in any phenotype that can be probed through a haploid genetic screen. Identification of such elements may not be feasible with RNA interference-based screens because they require that the target genomic sequences be transcribed. CRISPR-based technologies to screen for regulatory modules on a genome scale are still limited by the focused mutagenesis or transcriptional modulation of predetermined sequences in the genome [19–22]. However, focused CRISPR-based approaches would be powerful tools to precisely delineate any regulatory element found though our analysis.

While we found new regulatory elements in two central regulators of WNT signaling, we note that our current study is most likely under-powered to comprehensively detect all regulatory elements in the genome affecting the WNT pathway for several reasons. First, we used deep sequencing datasets from previous screens [3] that were designed to uncover protein coding genes involved in WNT signaling. The sequencing depth used to map insertions in these previous screens was sufficient to saturate the protein-coding genome, but is insufficient to interrogate the much larger non-coding genome. Second, the propensity of the retroviral-based mutagen used in this study to insert near TSSs, promoters and enhancers limited our search for regulatory elements to regions within and adjacent to genes. Our methodology could in principle be extended to identify regulatory elements located anywhere in the genome by using agents that integrate in a truly unbiased manner and then exhaustively mapping insertions in both the selected and unselected cell populations by sequencing at greater depth. Finally, because we assigned bins disregarding gene boundaries, our analysis may have missed regulatory elements in bins that overlapped with both an exon and an intron (such bins would have been excluded from the antisense intronic insertion enrichment analysis), and elements in bins that overlapped with features located both upstream and downstream of the TSS (such bins would have been excluded from the upstream insertion enrichment analysis). Reducing the size of the bins could ameliorate this problem, but at the expense of statistical power to determine the significance of GT insertion enrichment due to a reduction in GT insertions per bin and an increase in the multiple testing correction for a larger number of bins. Alternatively, computing GT insertions in intervals defined by the boundaries of genetic features such as introns or promoters (rather than bins of a predetermined size) could also help this issue, but would limit the analysis to annotated regions of the genome.

The analysis described in this work provides an untargeted and genome-scale method to identify both genes and regulatory elements involved in any biological process that can be probed by a haploid genetic screen. We hope that this bioinformatics analysis, available through Github (https://github.com/RohatgiLab/BAIMS-Pipeline), empowers other researchers to extract new insights about gene regulation from the growing body of insertional mutagenesis screen data.

## Acknowledgements

We thank members of the Rohatgi, Salzman and Carette labs for input on the project.

---

**S1 File. BAIMS output.**
Ranked lists of bins from the bin-based analyses. Each sheet of the Excel file contains a ranked list of bins determined by either the antisense intronic, upstream, or inactivating insertion enrichment analysis applied to either the WNT positive regulator low stringency or high stringency screen. The screen and type of bin-based analysis is indicated at the top of every sheet. The location of each bin in the human genome, the genes overlapping with the bin, and the FDR-corrected *p*-values generated by the bin-based analysis are specified. For each bin, the number of antisense intronic insertions, upstream insertions (sum of sense and antisense insertions), or inactivating insertions (sum of sense and antisense insertions for bins overlapping with a 5’UTR, CDS, or 3’UTR, or sense insertions only for bins overlapping exclusively with an intron) found within the bin in unsorted (control) and sorted cells are also indicated. The total number of insertions mapped in the unsorted cells and in the sorted cells are also shown.

**S2 Fig. Possible outcomes of GT insertions in different genetic features.**
(A) A GT consists of direct long terminal repeats (LTRs), a strong splice acceptor (SA), a reporter gene (mCherry) and a poly-adenylation (pA) sequence. A schematic of the 5’ end of a gene, including the promoter region, is also shown. (B) A GT can disrupt a gene by inserting into an exon in the sense orientation (with respect to the coding sequence of the gene), interrupting the coding sequence and causing premature transcriptional termination due to the pA sequence. (C) An antisense GT insertion into an exon interrupts the coding sequence of the gene and typically causes a frameshift mutation that leads to premature translational termination, producing a truncated protein. (D) When a GT integrates into an intron in the sense orientation, the SA causes the reporter gene and pA sequence to be spliced to the preceding exon, inevitably leading to premature transcriptional termination due to the pA sequence. (E) An antisense GT insertion in an intron will typically not disrupt a gene due to the directionality of the SA; however, it could interfere with regulatory elements or with transcripts present on the antisense strand. (F, G) GT insertions in the promoter region of a gene in either the sense or antisense orientation generally do not affect transcription of the downstream gene. However, they could potentially disrupt regulatory elements and alter transcription.

**S3 Fig. GT insertion patterns found in *TFAP4* and *DOT1L* in the WNT positive regulator low stringency and high stringency screens.**
(A) The histogram indicates the number and orientation of insertions mapped to *TFAP4* in the sorted cell populations from the WNT positive regulator low stringency and high stringency screens. See legend to Fig 3A for details. (B) The histogram indicates the number and orientation of insertions mapped to *DOT1L* (Chromosome 19, 2163750-2232749 bp) in unsorted cells and in the sorted cell populations from the WNT positive regulator low stringency and high stringency screens. The pattern of GT insertions seen in *DOT1L*, predominantly enriched for sense insertions in the first intron, differs from the observed enrichment for both sense and antisense insertions seen in the first intron of *TFAP4*.

**S4 Fig. GT insertion patterns found in *LRP6* in the WNT positive regulator low stringency and high stringency screens, and immunoblot analysis of LRP6.**
(A) The histogram indicates the number and orientation of insertions mapped to *LRP6* and to the region ~12.5 kbp upstream of the TSS in the sorted cell populations from the WNT positive regulator low stringency and high stringency screens. See legend to Fig 4A for details. (B) The histogram shows an expanded view of the 5’ end of *LRP6* and the region ~12.5 kbp upstream of the TSS (left of the vertical dotted line), with traces for GT insertions mapped in the sorted cell populations from the WNT positive regulator low stringency and high stringency screens. See legend to Fig 4B for details. (C) Immunoblot analysis of LRP6. The top and bottom parts of the same membrane were probed for LRP6 and ACTIN (loading control), respectively. The cell lines from which the samples were prepared and loaded in duplicate are indicated above the blots. Molecular weight standards in kDa are indicated to the left of each panel.

**S5 File. List of clonal cell lines containing GT insertions.**
The genomic sequences flanking GT insertion sites in the clonal cell lines used in this study are described. The first column (“Clone Name”) indicates the names of the clonal cell lines and the second column (“Genomic sequence flanking GT”) indicates the genomic sequences 5’ and 3’ from the GT insertion site (relative to the sense orientation of the GT as described in S2 Fig).

